# Tools for the genetic manipulation of *Herpetomonas muscarum*

**DOI:** 10.1101/2020.01.03.894162

**Authors:** Megan A. Sloan, Petros Ligoxygakis

## Abstract

Trypanosomatid parasites are causative agents of important human and animal diseases such as sleeping sickness and leishmaniasis. Most trypanosomatids are transmitted to their mammalian hosts by insects, often belonging to Diptera (or true flies). With resistance to both vector-targeted pesticides and trypanocidal drugs being reported, there is a need for novel transmission blocking strategies to be developed. Studies using the blood-feeding vectors themselves are not broadly accessible, as such, new model systems are being developed to unpick insect-trypanosmatids interactions. One such case is the interactions between the model dipteran *Drosophila melanogaster* and its natural trypanosomatid *Herpetomonas muscarum*. Our previous work has found that much of the transcriptomic changes triggered in *H. muscarum* after ingestion by *Drosophila* reflect what is known for disease-causing trypanosomatids. Here we describe a set of tools to genetically manipulate the parasite and therefore create a truly tractable insect-parasite interaction system from both sides of this association. These include transgenic fluorescently tagged parasites to follow infection dynamics in the fly gut as well as iterations of plasmids that can be used for generating knock-in and knock-out strains. The tools presented in this short report will facilitate further characterisation of trypanosomatid establishment in a model dipteran.

## Introduction

Neglected Tropical Diseases are the most common diseases of the world’s poorest people. Many are caused by trypanosomatid, single-celled protozoans, parasites which are transmitted to humans via insects belonging to the order of Diptera (also known as true flies). These flies (including tsetse, sand flies and black flies) are difficult to study in the lab and so the prospect of rapid progress in the basic biology of fly-parasite interaction is bleak. However, a model dipteran species with an extensive “tool-box” is the fruit fly *Drosophila melanogaster* which harbours the trypanosomatid *Herpetomonas muscarum* in wild populations. We have begun to use this system to identify evolutionarily conserved aspects of dipteran-trypanosomatid interactions^1–3^. One of the strengths of this model is the plethora of knowledge, tools and resources available for the study of *D. melanogaster*. However, despite extensive work in other trypanosomatids available^4–9^, there are no published examples of *H. muscarum* transgenic lines. As such, we developed genetic tools for use in this understudied trypanosomatid based on those described for *Leishmania*^8,9^.

## Results and Discussion

### Producing transgenic *Herpetomonas muscarum*

We constructed novel plasmids based on those developed in Dean *et al*. 2015. These plasmids contain the pLENT ‘backbone’ (generous gift from Dr Eva Gluenz, Oxford) enabling replication and cloning in *Escherichia coli*, and an expression ‘cassette’ flanked by the whole intergenic regions of *H. muscarum* beta-tubulin gene (HMUS00495500). The whole intergenic regions were used as the specific sequences required for regulation of gene expression in these regions are not known in *H. muscarum*. The expression cassette contains the open reading frame for a fluorescent protein e.g. tdTomato, the intergenic region between phosphoglycerate kinases A and B (PGKAB) and an antibiotic resistance gene to enable selection (Figure S1). Several iterations of pMS003 have been produced to allow the expression of several different fluorescent proteins under the selection of four different antibiotics (Table S1).

Drug resistant cells, expressing tdTomato were successfully produced after electroporation with two pulses of 1500V for 100μs (with a 200ms gap between pulses) in the presence of plasmid which had been linearized by restriction digest using BSPDI (Figure 1). The minimum mass of plasmid required to produce transgenic *H. muscarum* using the pMS003 vector was found to be between 10-20μg of linearized plasmid DNA – approximately 1.75-3.5 pmol of DNA. We were unable to purify plasmid from the transgenic cell lines, as such the expression cassette appears to have integrated into the genomic DNA.

**Figure 1 –.**
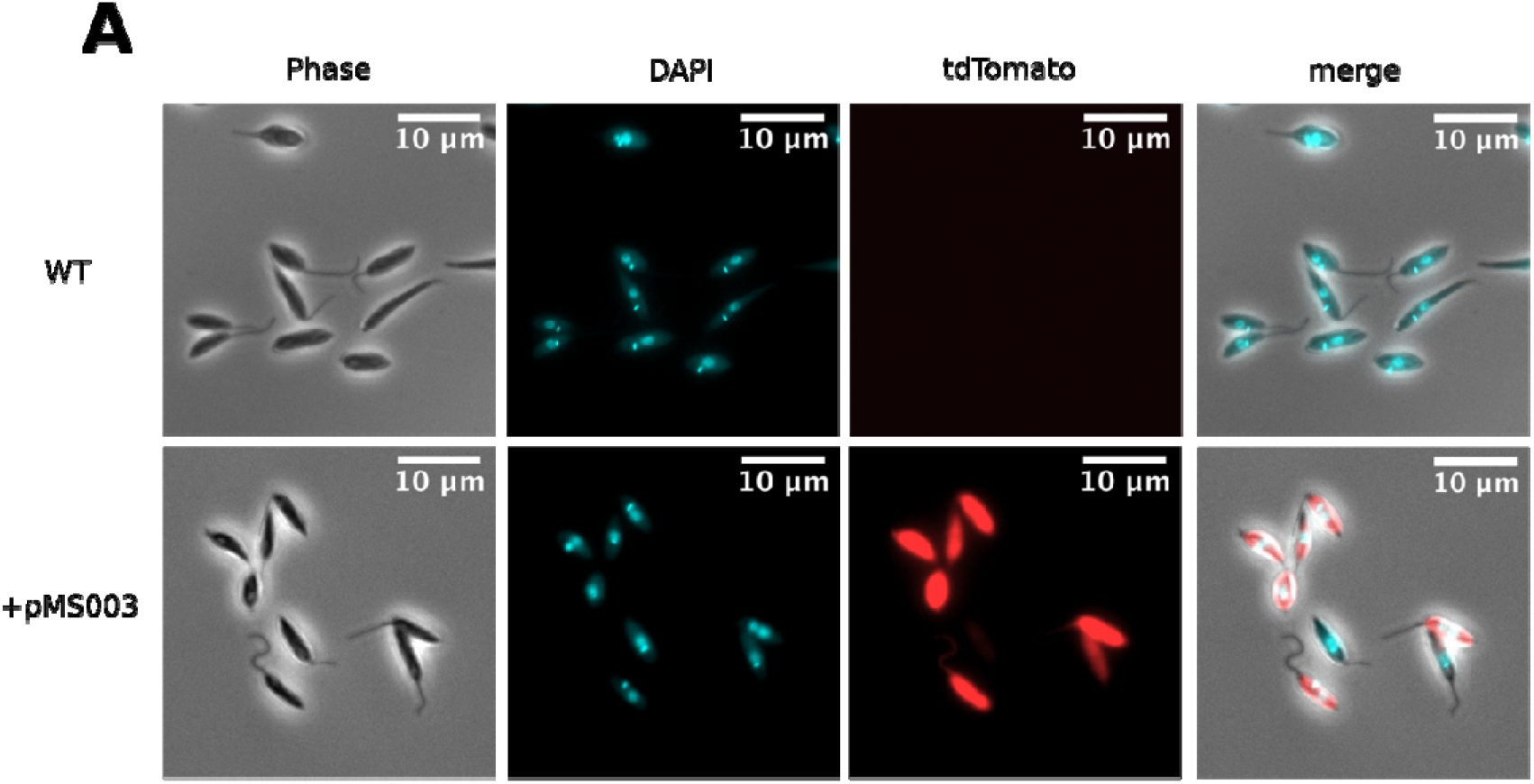
Light microscopy of *H. muscarum* transfected with pMS003 and wild type *H. muscarum*

### Following *H. muscarum* infection in *D. melanogaster*

Having produced a tdTomato^+^ cell line we sought to use this line to follow the trypanosomatids after ingestion. Analysis of the growth curves shows the transgenic cells grow slower than their wild type counterparts – which is typical of trypanosomatids grown under drug selection (Figure 2). However, we did not find any significant difference in parasite loads during *D. melanogaster* infection between the tdTomato^+^ and wild type lines (Figure).

**Figure 2 –.**
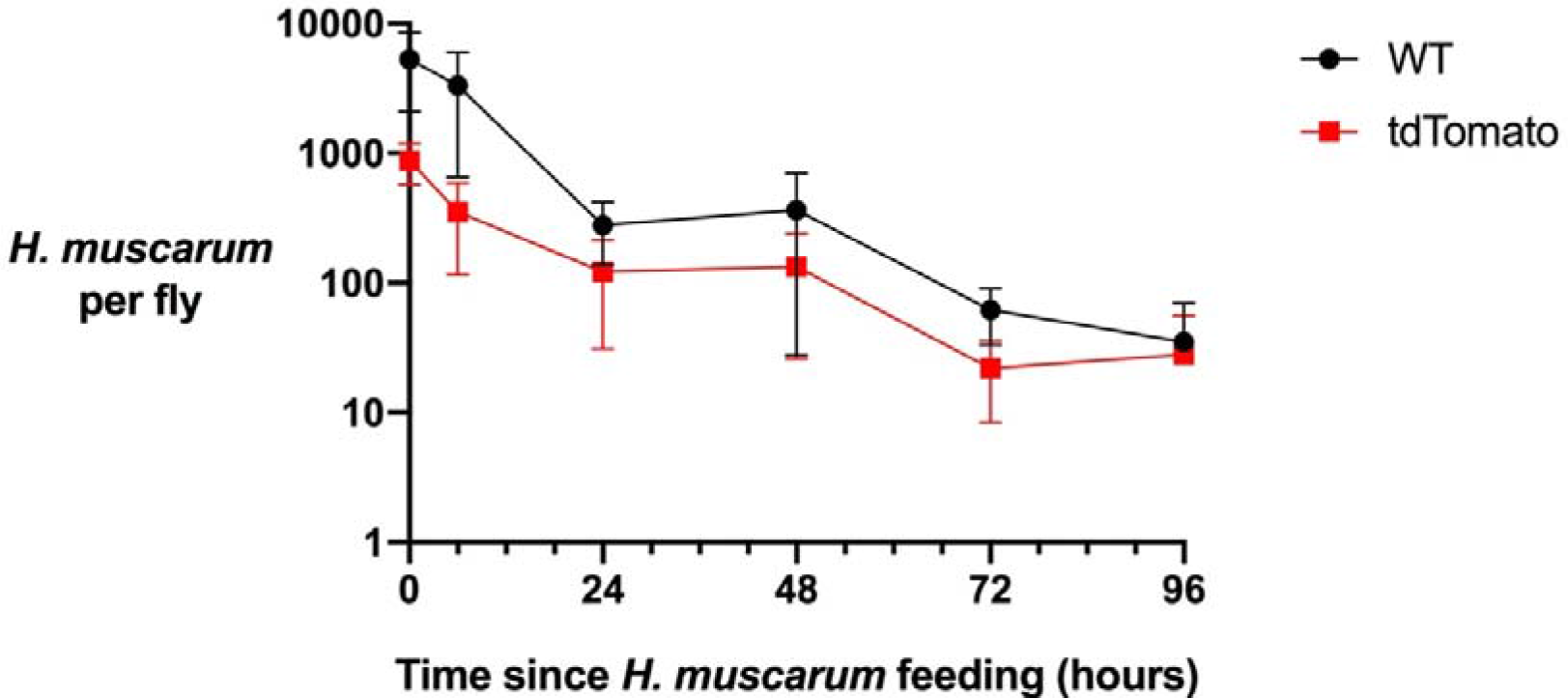
*Herpetomonas muscarum* load in *Drosophila melanogaster* after two hours feeding. The average number of *H. muscarum* per fly for three biological repeat infections and error bars show the standard error of the mean. WT – wild type, tdTomato – *H. muscarum* expressing the tdTomato fluorescence protein.

Imaging of dissected guts from flies immediately after following feeding showed that *H. muscarum* cells enter the crop and midgut of the flies in large numbers (Figure 3). At six hours post feeding there were still many trypanosomatids in the crop of the flies, however *H. muscarum* were no longer dispersed throughout the crop but appeared restricted to a smaller area close to the crop wall (Figure 4). The majority of *H. muscarum* cells in these tightly packed clusters are rounded – a phenotype commonly associated with stress in trypanosomatids. During imaging of live (dissected, unfixed) gut preparations these cells have beating flagella and as such appear to be alive. In contrast to immediately after feeding, there were very few cells visible in the midgut at six hours, often in small groups of cells (Figure).

**Figure 3 –.**
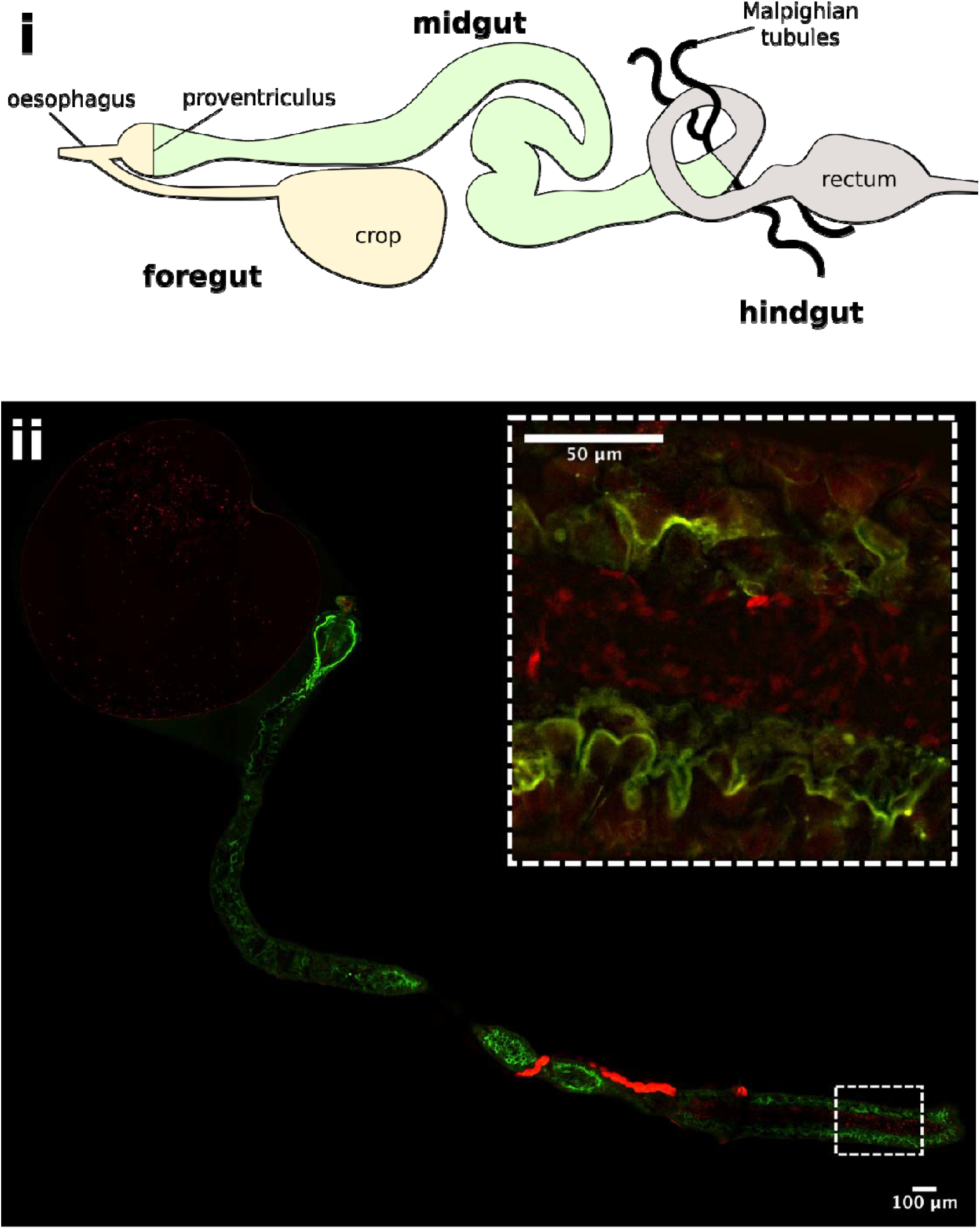
*Herpetomonas muscarum* in the fly digestive tract immediately after feeding. i - Schematic of the *D. melanogaster* digestive tract. ii – The foregut and midgut of *D. melanogaster* two hours after feeding with tdTomato expressing *H. muscarum*. The flies used for this work express a myosin-GFP fusion protein to allow the gut epithelial border to be visualised. *H. muscarum* can be seen in the crop and the midgut (inset) of the fly.

**Figure 4 –.**
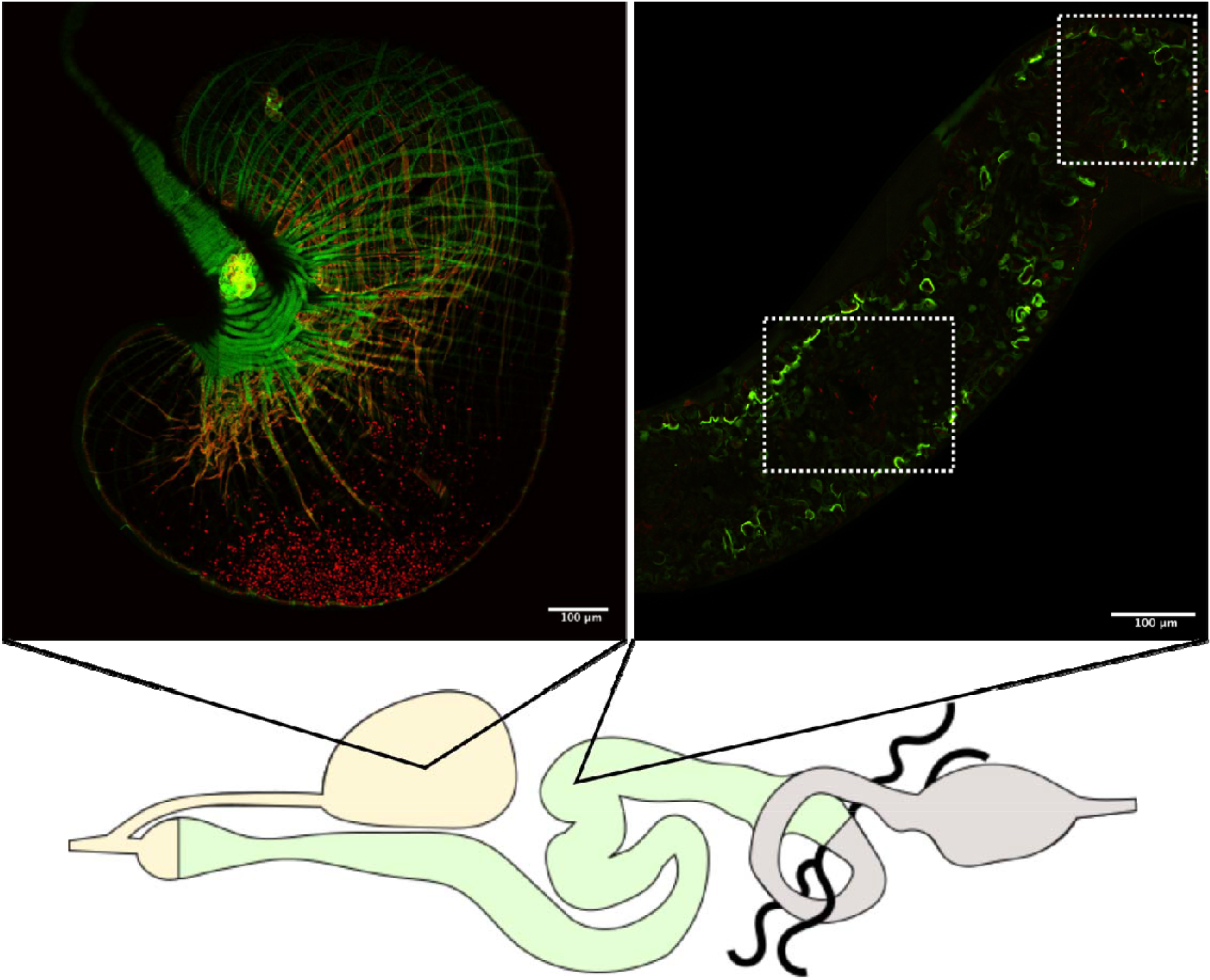
*Herpetomonas muscarum* are found in the fly crop and midgut 6 hours after feeding. *H. muscarum* cells expressing tdTomato were fed to *D. melanogaster* expressing a myosin-GFP fusion protein to allow the gut epithelial border to be visualised. *H. muscarum* can be seen mainly in the crop, clustered, close to the epithelial layer. Small numbers of *H. muscarum* cells can be seen in the midgut (highlighted by the white-dashed boxes).

## Conclusions

We have produced a suite of plasmids allowing exogenous expression of proteins in *H. muscarum*. We have demonstrated the use of these plasmids to express fluorescent protein tdTomato. To our knowledge this is the first known genetic manipulation of *H. muscarum*. These lines have been used for the localisation of *H. muscarum* cells in the *D. melanogaster* digestive tract, specifically the crop and midgut, following ingestion.

These plasmids are modular and will allow, after two rounds of PCR, the tagging and knockout genes of interest in *H. muscarum* as described in Dean *et al*. 2005^8^. Use of multiple expression cassettes with different selection drugs will also allow the production of double “mutants”. As such, the plasmids reported herein will be a useful tool for the study of *H. muscarum*, both in the context of the *Drosophila-Herpetomonas* model as well as for the almost 200 species of Diptera that can be infected by *Herpetomonas muscarum*^10–12^.

## Materials and methods

### Production of pMS003

The components of the expression cassette were amplified by PCR so that they contained an 18bp overlap with the adjacent fragment in the cassette (for primer sequences see Table S2). This allowed the fragments to be joined to each other, and the pLENT backbone using the NEBuilder Assembly kit. After assembly reaction, 2μL of the resulting mix was transformed into chemically competent *Escherichia coli* (High efficiency DH5-alpha, New England Biolabs) according to manufacturer’s instructions and cultured for 16 hours on Luria broth (LB) agar plates with 100μg ml^−1^ ampicillin. Individual clones were then assayed by colony PCR to check for the presence of the fluorescent protein (or gene of interest). ‘Positive’ colonies were used to inoculate 50ml of LB with 100μg ml^−1^ ampicillin which was cultured, shaking at 37°C for 16 hours. Plasmid was purified from the overnight culture using the Qiagen Midi Prep kit (cat # 12143) and the expression cassette sequenced prior to use for transfection of *H. muscarum*.

### *H. muscarum* transfection

Cells were taken from log phase *in vitro* culture and pelleted by centrifugation at 800xg for 3 minutes. The supernatant was removed and the cells resuspended in either; Zimmerman’s (Zimmermann and Vienken, 1982) electroporation buffer to a concentration of 8E7 cells ml-1. For each electroporation 200μL of cell suspension was used. Cells were added to 0.4cm gap electroporation cassettes (Biorad) with plasmid DNA and electroporated. After electroporation cells were added to pre-warmed media and incubated at 28^º^C for 12 hours before the addition of the selection drug.

### DNA extraction from *H. muscarum*

*H. muscarum* cell suspension used for feeding was retained and had the gDNA extracted using the Norgen Biotek Genomic DNA Isolation kit (Cat# 24700) according to the manufacturer’s instructions.

### Light microscopy of *H. muscarum*

Cells were pelleted and washed three times in sterile phosphate buffered saline (PBS) before being resuspended in 100μL of PBS per half slide. The cell suspension was pipetted onto a glass slide, whose borders were marked out with a hydrophobic barrier (DAKO pen, Agilent Technologies Ltd.) and the cells allowed to settle for 10 minutes. The slides were moved to a dark, humidified chamber for subsequent steps. Cells were then fixed by adding 100μL of 4% paraformaldehyde to the slide which was left at room temperature for 10 minutes. The slides were rinsed by three successive immersions in clean PBS then permeabilised by incubation in ice cold methanol for 30 minutes. The cells were then re-hydrated in PBS for 5 mins. Nucleic acids were stained with DAPI (4□,6-diamidino-2-phenylindole), a blue-fluorescent DNA stain that exhibits enhanced fluorescence when bound to double stranded DNA, by covering the cells with 100μL of 1μg ml-1 DAPI solution in PBS for 10 mins. The slides were rinsed by three successive immersions in clean PBS and sealed with a coverslip and nail varnish. All slides were stored at 4°C in the dark. Cells were imaged using the Leica DM5500 B and Andor Neo sCMOS camera system.

### Infection of *D. melanogaster*

Prior to infection seven-day old (from eclosion) adult D. melanogaster were taken off conventional fly food and given access only to water for 12 hours prior to infection. Log-phase H. muscarum cultures were pelleted at 800xg for 3 minutes, the supernatant removed and then resuspended in standard H. muscarum culture media with 10% sucrose to a cell density of 2×107 cells ml-1. Half a millilitre of this cell suspension was pipetted onto two 22mm filter paper discs (Whatman™) placed in the base of a clean, dry fly vial. Drosophila melanogaster were added to the vial and incubated under standard conditions for 2-4 hours. After this, the flies were transferred back onto conventional fly food for the remainder of the experiment.

### Light microscopy of infected *D. melanogaster* digestive tracts

Flies were anesthetised using CO2 prior to dissection. Immediately post-dissection the fly guts were fixed in 200μL of 4% paraformaldehyde in PBS for 1 hour. The guts were then rinsed three times in 500μL of PBS, on the bench at room temperature, for 15 minutes per wash. The guts were then mounted on a glass slide in a drop of halocarbon oil (VWR) and sealed with a coverslip and nail varnish. The slides were then stored at 4°C and imaged with 48 hours. Mounted digestive tracts were imaged with the Olympus FV3000 confocal system (fluorescence images) and the Zeiss Imager.Z2 microscope with Hamamatsu Orca Flash4.0 camera (phase contrast images). ImageJ software was used for image handling and file-type conversions.

## Acknowledgements

We would like to thank Dr. Eva Gluenz, Prof. Keith Gull and members of their labs for their advice and guidance during this work.

## Supplementary Figures

**Figure S1 –.**
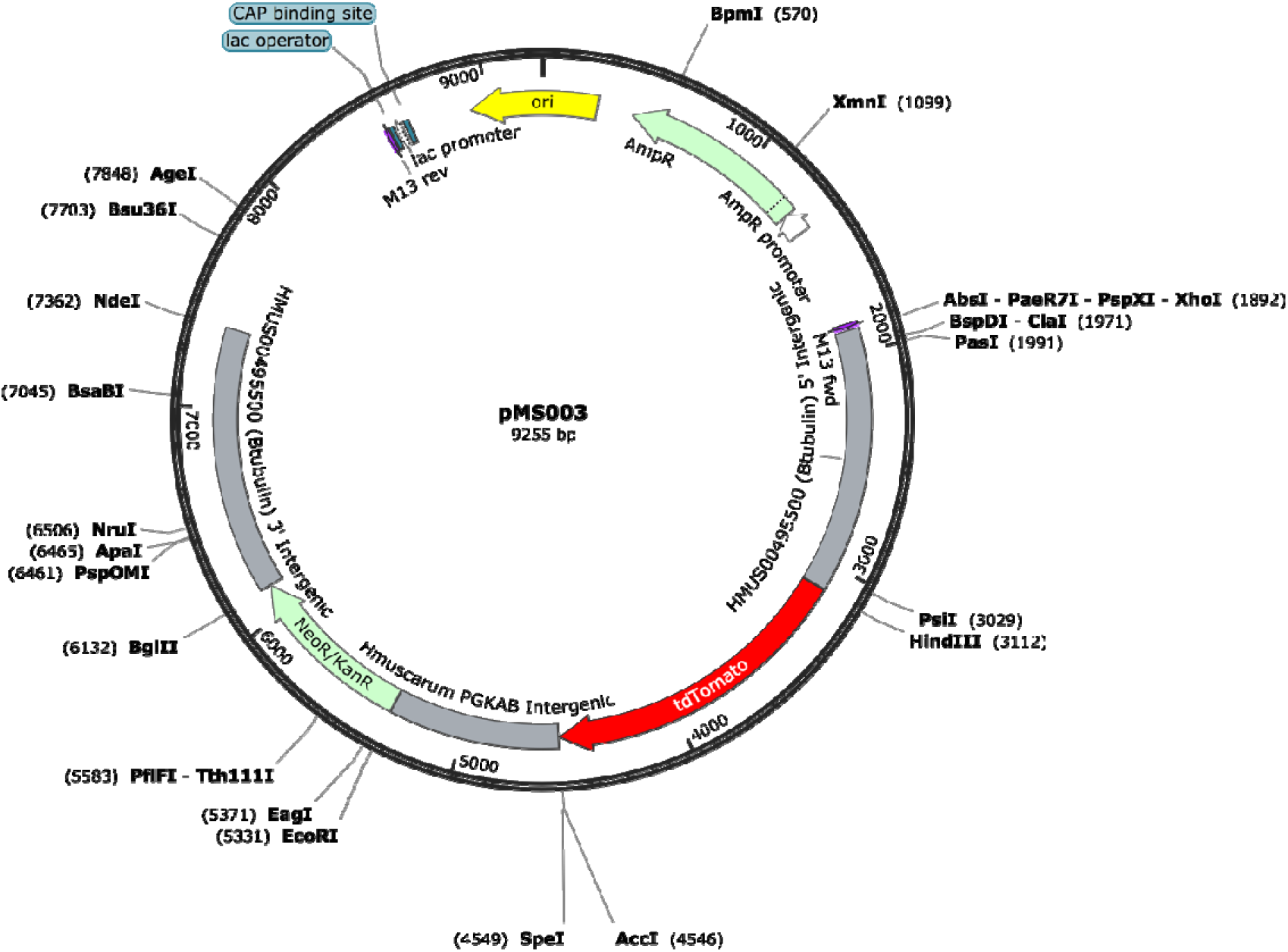
Plasmid map of pMS003 showing unique 6+ cutter restriction enzyme sites. This image was produced using SnapGene software.

## Supplementary Tables

**Table S1 –.**
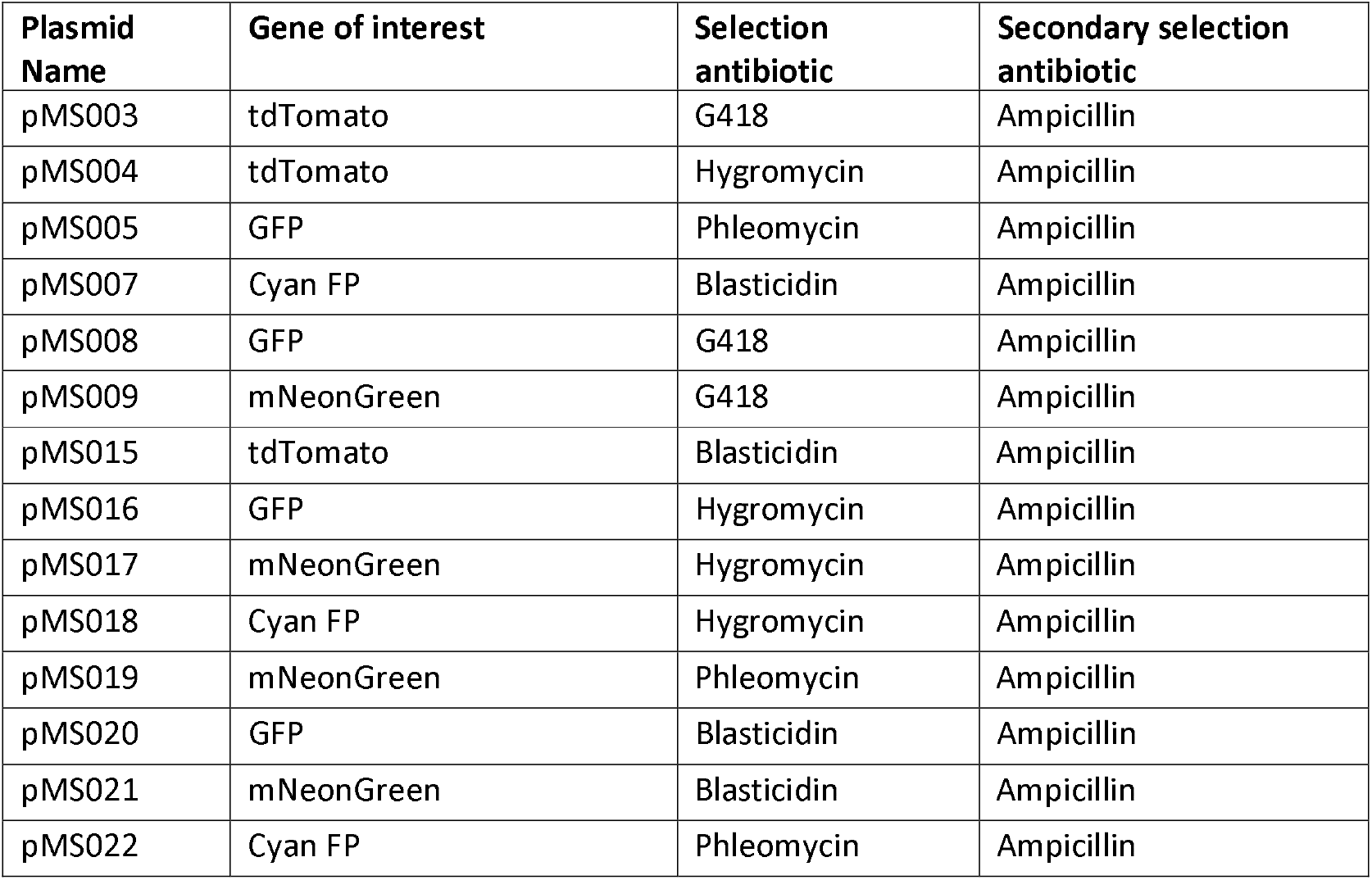
A list of plasmids produced in this work allowing the expression of a fluorescent protein in *Herpetomonas muscarum*

**Table S2 –.**
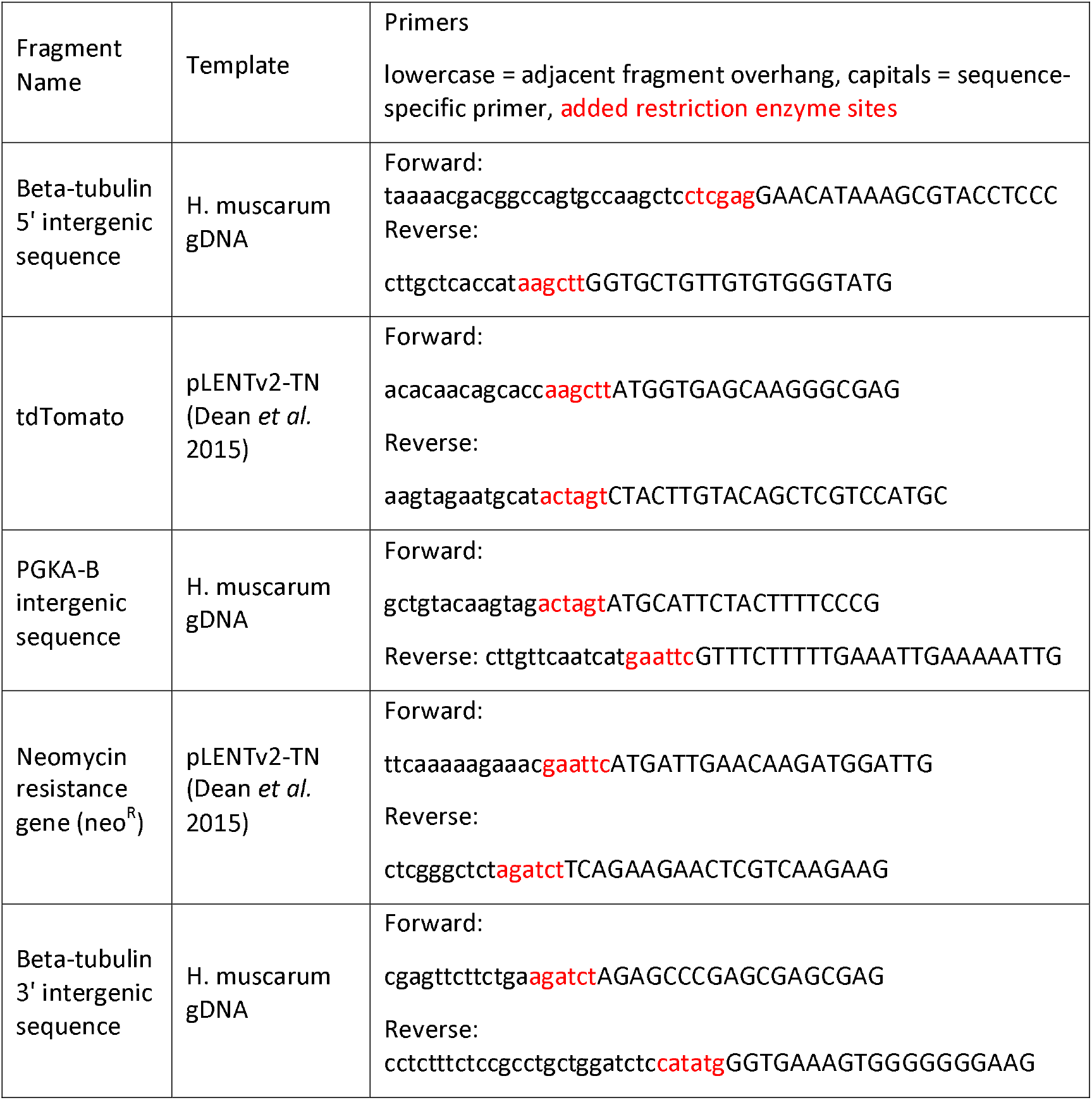
Primers for construction of novel expression vector pMS003. Restriction sites (red): CTCGAG = XhoI, AAGCTT=HindIII, ACTAGT=SpeI, GAATTC=EcoRI, CATATG=NdeI.

## Supplementary data files

**The plasmid sequence for pMS003.** This plasmid allows the expression of the fluorescent protein tdTomato in H. muscarum under selection of the antibiotic neomycin. It may also be used to knock out or tag the C-terminus of *H. muscarum* genes of interest.

